# Widespread transposon co-option in the *Caenorhabditis* germline regulatory network

**DOI:** 10.1101/2022.01.21.477252

**Authors:** Francesco Carelli, Chiara Cerrato, Yan Dong, Alex Appert, Abby Dernburg, Julie Ahringer

## Abstract

The movement of selfish DNA elements can lead to widespread genomic alterations with potential to create novel functions. Here we show that transposon expansions in *Caenorhabditis* nematodes led to extensive rewiring of germline transcriptional regulation. We find that about one third of *C. elegans* germline-specific promoters have been co-opted from two related Miniature Inverted Repeat Transposable Elements (MITEs), CERP2 and CELE2. The promoters are regulated by HIM-17, a THAP domain-containing transcription factor related to a transposase. Expansion of CERP2 occurred prior to radiation of the *Caenorhabditis* genus, as did fixation of mutations in HIM-17 through positive selection, whereas CELE2 expanded only in *C. elegans*. Through comparative analyses in *C. briggsae*, we find evolutionary conservation of most CERP2 co-opted promoters, but a substantial fraction of events are species specific. Our work reveals the emergence of a novel transcriptional network driven by TE co-option with a major impact on regulatory evolution.

## Introduction

Cis-regulatory elements play fundamental roles in gene expression yet can undergo remarkably rapid evolutionary turnover (1–3). Transposable elements (TEs) are a potential source of novel regulatory elements as they harbor regulatory sequences recognized by the host machinery. If moved to an appropriate location, such sequences may affect the expression of host genes, and clear evidence for co-option of some TE insertions into host regulatory networks has been documented (4, 5). It has been suggested that the amplification of a TE family could lead to the co-option of many TEs, dramatically changing whole regulatory networks (e.g. (6–9)), but the demonstration of such events is a challenging task. Ancient co-options would likely be masked by mutations that obscure their repetitive origin, while the functional relevance of recent co-options can be hard to determine on a large scale. As a result, there is limited functional evidence *in vivo* to support widespread or concerted transcriptional rewiring. Here we show through genomic and functional analyses in *Caenorhabditis* that two independent TE expansions gave rise to promoters that control the expression of a large fraction of germline-specific genes.

## Results and Discussion

To investigate transcription regulation in the *C. elegans* germline, we first identified germline-specific accessible chromatin sites (N=2316) based on the presence of a strong ATAC-seq signal in wild-type young adults but not in *glp-1* mutants lacking a germline (10)(Fig S1A). Using chromatin-associated RNA-seq patterns to link open chromatin regions to annotated genes, we then classified 782 sites as germline-specific promoters (Fig 1A; Table S1; see Methods). Sequence analysis of these promoters revealed the enrichment of two motifs (m1 and m2; Fig 1B) that do not share significant similarity with other eukaryotic regulatory motifs, but were previously identified upstream of genes with germline expression (11). We found that an m1m2 pair is present in 36.3% (284/782) of all germline-specific promoters. These motifs were more commonly found in divergent orientation (29.8% of promoters), while the other 6% showed a tandem arrangement (m2m1, Fig 1C). Promoters containing m1m2 motifs were also found upstream of 177 genes expressed in both germline and soma, which predominantly show ubiquitous accessibility by ATAC-seq (Fig S1B). Genes associated with m1m2-containing promoters are more highly expressed than other germline genes, and their promoters show greater accessibility in primordial germ cells (PGCs) as well as in late larvae, which contain many germline cells (Fig S1C,D).

**Fig. 1.**
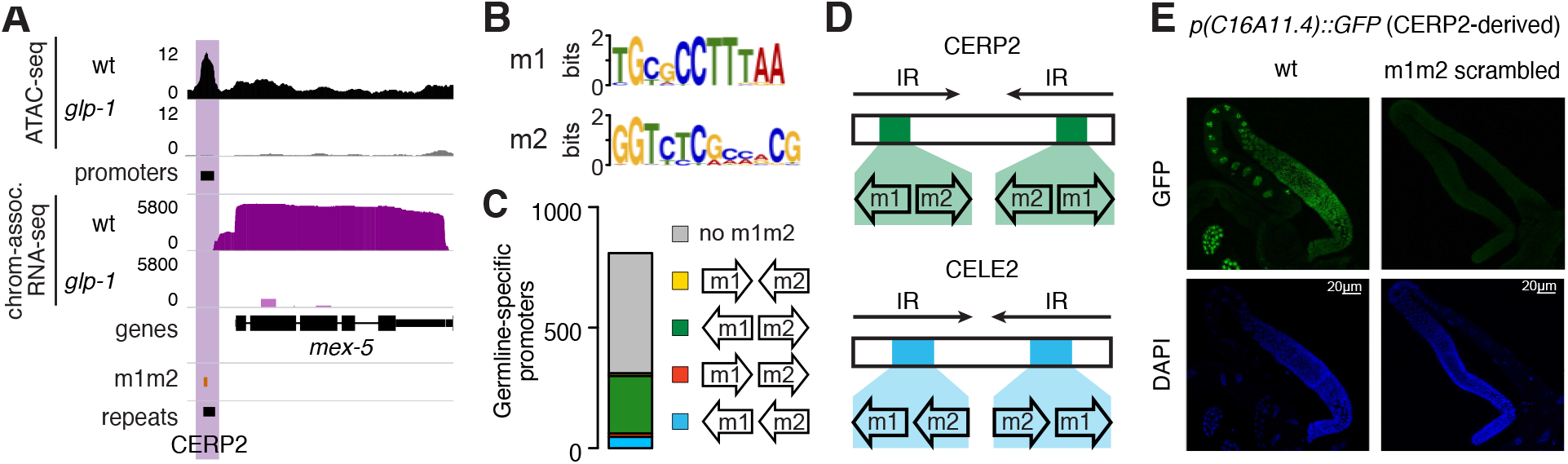
TE enrichment at germline-specific elements in *C. elegans*. (**A**) Example of germline-specific (purple) promoter in *C. elegans*. (**B**) Sequence logos of the m1 and m2 motifs. (**C**) Number of m1m2 pairs overlapping germline-specific promoters, color-coded based on their relative orientation. (**D**) Location of m1m2 pairs in CERP2 and CELE2 consensus. IR: inverted repeats. (**E**) GFP and DAPI signals from CERP2-derived wt *p*(*C16A11.4*)::*his-58::gfp* and m1m2-scrambled *p*(*C16A11.4*)::*his-58::gfp* in adult gonads (scale bar 20 μm).

While m1m2 pairs were strongly associated with germline promoters, many additional copies of these motifs were also found in non-accessible regions of the *C. elegans* genome in both divergent (n=1458) and tandem (n=2566) orientations, and were characterised by distinct m1-m2 spacing distributions (Fig S1F). These predominantly corresponded to the positions of CERP2 and CELE2 elements, respectively, which are also the most highly enriched repeat classes at germline-specific promoters (Fig S1E,G; Table S1). These comprise two families of Miniature Inverted Repeat Transposable Elements (MITEs), small, non-autonomous elements derived from autonomous DNA transposons (12, 13). The inverted repeats of both elements contain m1 and m2 motifs, oriented divergently in CERP2 and tandemly in CELE2 (Fig 1D). The similar structure, and the presence of the motif pair suggests an evolutionary relationship between CERP2 and CELE2, yet their origin is unclear since they do not share any similarity with annotated autonomous transposons. We found that m1m2 promoters matched CERP2 and CELE2 consensus sequences with similar identity scores to non-promoter m1m2 pairs, supporting derivation of m1m2 promoters from MITE elements (Fig S1H). Both promoter- and non-promoter-associated copies of CERP2, and to some extent the CELE2 family, also contain a region of 10-bp periodic TT bias. This feature was recently shown to be associated with nucleosome positioning in *C. elegans* germline promoters (14)(Fig S1I), and may have facilitated the co-option of these MITEs by creating a chromatin environment which facilitates transcription in this tissue. These results show that a large fraction of *C. elegans* germline-specific promoters are derived from CERP2 and CELE2 MITEs.

To understand the relevance of the m1 and m2 motifs in MITE-derived promoters for germline transcription, we introduced wild-type and mutant transgenes into *C. elegans*. CELE2 and CERP2 derived promoters with wild-type m1m2 sequences drove germline-specific expression of a histone-GFP reporter (Fig 1E, Fig S1J). We found that both motifs were required for promoter activity, as GFP was not detectable after scrambling m1 or m2 (Fig 1E, Fig S1J). In addition, scrambling the motif sequences in the endogenous CERP2-associated T05F1.2 promoter using CRISPR-Cas9 editing reduced expression by 5.9-fold (Fig S1K). These results strongly support the idea that CERP2 and CELE2 elements were co-opted as germline-specific promoters, and show that the m1 and m2 motifs are required for their regulatory activity.

To identify a potential transcription factor that might regulate these co-opted promoters, we analysed modENCODE transcription factor binding data (15) for enrichment at co-opted versus non-co-opted germline promoters. We found that HIM-17 showed the highest enrichment (>7.6-fold, Fig S2A). HIM-17 is a germline chromatin-associated factor important for meiosis and germline organization (16, 17). It has six THAP domains, putative DNA binding domains shared by P-element family transposases (18).

As the HIM-17 ChIP-seq modENCODE data were from a mutant background, we generated new HIM-17 ChIP-seq data from wild-type adults, which identified 3539 HIM-17 peaks (Fig 2A, Table S1). HIM-17 binding was strongly associated with m1m2 motifs; all but one of the 284 co-opted germline-specific promoters were associated with a HIM-17 peak, as were 80.8% of non-germline specific m1m2-containing promoters (Fig 2B). HIM-17 peaks were also associated with non-promoter m1m2 pairs, including sites in closed chromatin environments (Fig 2C, Fig S2B). The m1m2 pair is the likely determinant of HIM-17 binding, as HIM-17 enrichment at a co-opted promoter was abolished when either m1 or m2 was mutated (Fig S2C).

**Fig. 2.**
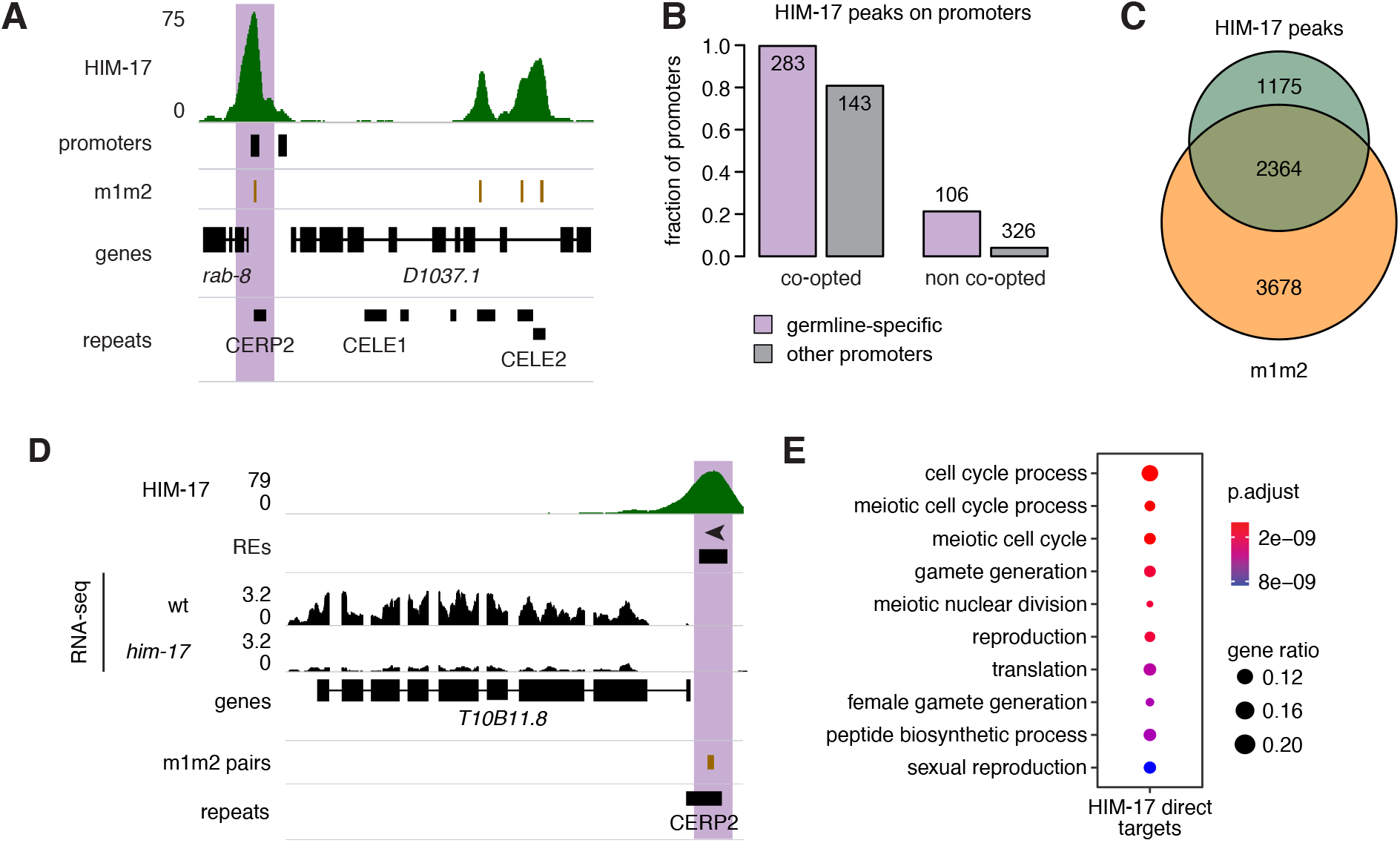
HIM-17 binds and regulates co-opted MITEs. (**A**) Example of HIM-17 ChIP-seq binding profile. (**B**) Fraction of promoters overlapped by HIM-17 peaks. (**C**) Overlap between HIM-17 peaks and m1m2 pairs. (**D**) Example of a gene downregulated specifically in *him-17* mutants. (**E**) GO terms enrichment of HIM-17 downregulated direct targets.

To determine whether HIM-17 plays a role in the expression of co-opted promoters, we analyzed gene expression in the strong loss-of-function mutant *him-17*(*me24*). Comparison of RNA-seq data between mutant and wild-type animals indicated that HIM-17 acts as a transcriptional activator, because only genes showing lower expression in the mutant were significantly associated with HIM-17 binding at their promoters (Fig 2D, Fig S2D, Table S1). Based on HIM-17 binding, we defined 304 genes as direct targets (Table S1). Gene ontology analysis revealed a strong enrichment for genes affecting meiosis and reproduction among direct targets of HIM-17, in line with meiotic defects documented in *him-17* mutants (16)(Fig 2E). Notably, among the genes strongly downregulated in *him-17*(*me24*) mutants were *him-5* and *rec-1* (Fig S2E), two paralogs that promote double-strand break (DSB) formation during meiosis (19). Downregulation of these HIM-17 targets likely accounts for the strong reduction in DSBs and the resulting High incidence of males (Him) phenotype associated with mutations in *him-17* (16, 20)(Fig S2E). The large number of genes regulated by HIM-17 also explains the pleiotropic effects of such mutations on meiosis and other germline processes (16, 17, 20–22). These findings indicate that HIM-17 acts as a transcriptional activator that regulates genes whose promoters were co-opted from MITEs.

To gain further insights into CERP2 and CELE2 co-option and their regulation by HIM-17, we investigated their evolution through comparative analyses in nematodes. We first sought to determine the timing of the co-option process by dating the TE expansion events. CERP2 elements were abundant in the genomes of all *Caenorhabditis* species we analyzed, but not in other nematodes (Fig 3A). In contrast, CELE2 elements were detected only in *C. elegans*, suggesting a recent, species-specific expansion of this repeat family (Fig 3B). The earlier expansion of CERP2 is also reflected in the higher proportion of truncated CERP2 copies compared to CELE2 in *C. elegans* (Fig S3A). In addition, we detected a high number of tandem m2m1 pairs in *C. becei* and *C. monodelphis* with distinct spacing of the m1 and m2 sequences relative to CELE2, suggesting that other related TEs likely underwent expansion in these *Caenorhabditis* species (Fig S3B). These data indicate that the CERP2 and CELE2 expansions took place at different times in the *Caenorhabditis* clade, seeding thousands of m1m2 sequences and generating a large reservoir of potential regulatory elements.

**Fig. 3.**
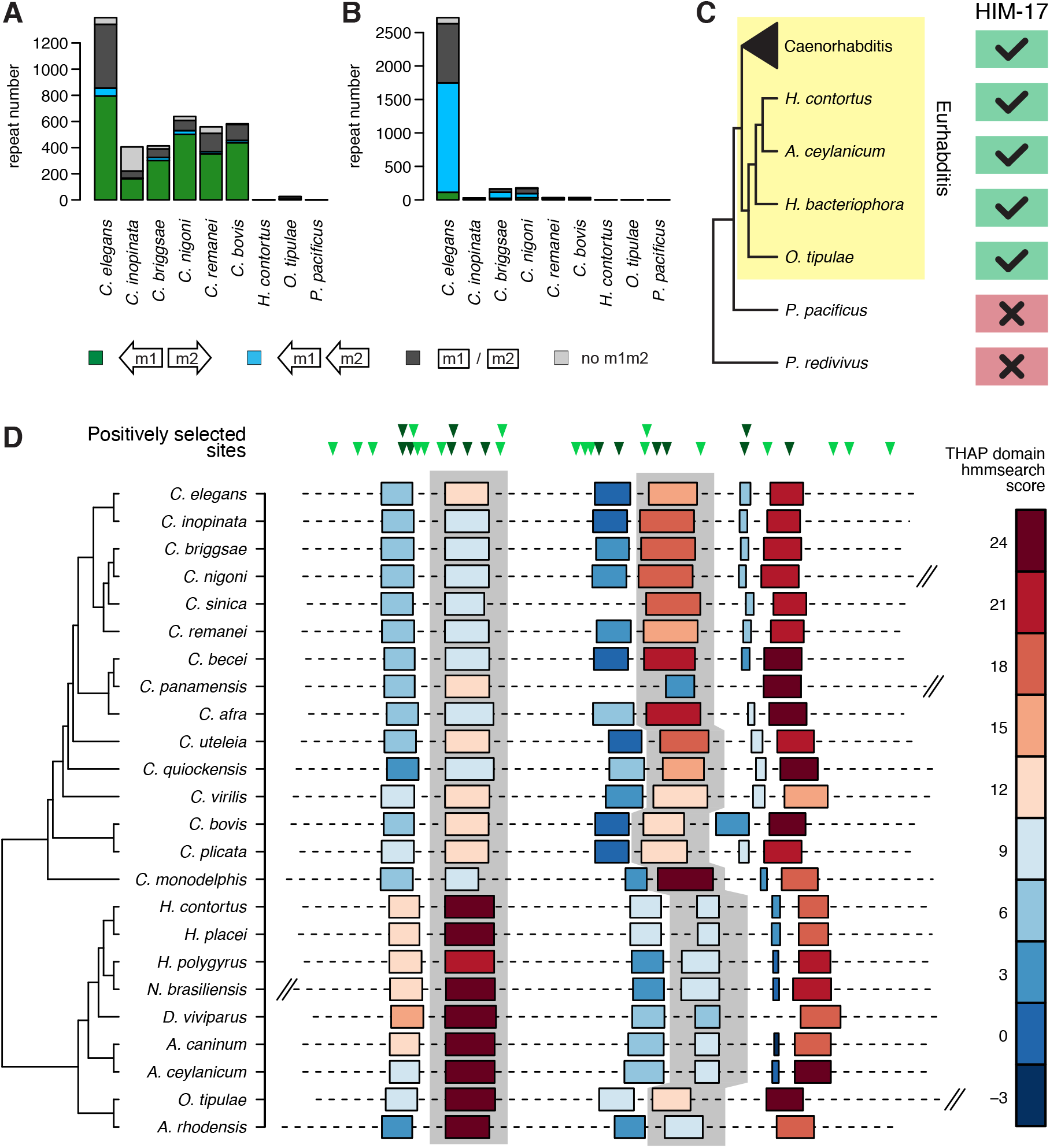
Evolution of m1m2 pairs and their binding factors in nematodes. (**A,B**) Number of CERP2 (A) and CELE2 (B) elements annotated in the genomes of different nematode species, and fraction of divergent_m1m2 pairs, tandem_m2m1 pairs, and other m1 and or m2 motifs overlapped. (**C**) Evolutionary conservation of *him-17*. (**D**) Top: location of sites under positive selection with respect to the *C. elegans* HIM-17 protein. In dark, sites located in a THAP domain. Bottom: location of THAP domains in HIM-17 orthologs. Color code reflects their similarity to the canonical THAP domain (based on the hmmsearch score). Second and fourth THAP domains are highlighted in grey. Protein length is drawn to scale, and truncated for longer orthologs.

HIM-17 predates the *Caenorhabditis*-specific expansions of CERP2 and CELE2, as orthologs could be identified not only in *Caenorhabditis* genomes, but also in in other Eurhabditis nematodes, with the exception of *Diploscapter* species (Fig 3C, Fig S3C,D). In light of the regulation of m1m2-associated promoters by HIM-17, we speculated that HIM-17 sequence might have undergone changes in line with the timing of the *Caenorhabditis* CERP2 expansion. Evolutionary analyses indicate that *him-17* underwent positive selection prior to divergence of the *Caenorhabditis* genus (branch-site test, P = 0.0007), as did expansion of the CERP2 sequence. 14/34 of the sites under positive selection are located within its 6 THAP domains (Fig 3D), which are related to the DNA-binding domain of the *Drosophila* P-element transposase (18) and conserved in almost all HIM-17 orthologs. Moreover, compared to the sister Strongylida clade, the fourth THAP domain in all *Caenorhabditis* species is more similar to the Pfam THAP consensus, while the second THAP domain is more divergent (Fig 3D). These conserved changes in putative DNA binding domains occurred in parallel with the CERP2 expansion, and we speculate that they may have enhanced HIM-17 recognition of the MITE-derived m1m2 motifs.

A large fraction of co-opted CERP2 sequences showed evidence of evolutionary conservation, as indicated by peaks of phyloP scores, a measure of sequence conservation across multiple species (Fig 4A). To directly evaluate and quantify whether co-option events in the *Caenorhabditis* genus have given rise to shared and/or lineage-specific regulatory elements, we analysed germline promoters in *C. briggsae*, which diverged from *C. elegans* ~20 million years ago (23). As we did for *C. elegans*, we identified *C. briggsae* germline specific promoters by generating ATAC-seq and nuclear RNA-seq data from wild type and a germline-less *C. briggsae glp-1* temperature-sensitive mutant that we generated using CRISPR editing (see Methods; Fig S4A).

**Fig. 4.**
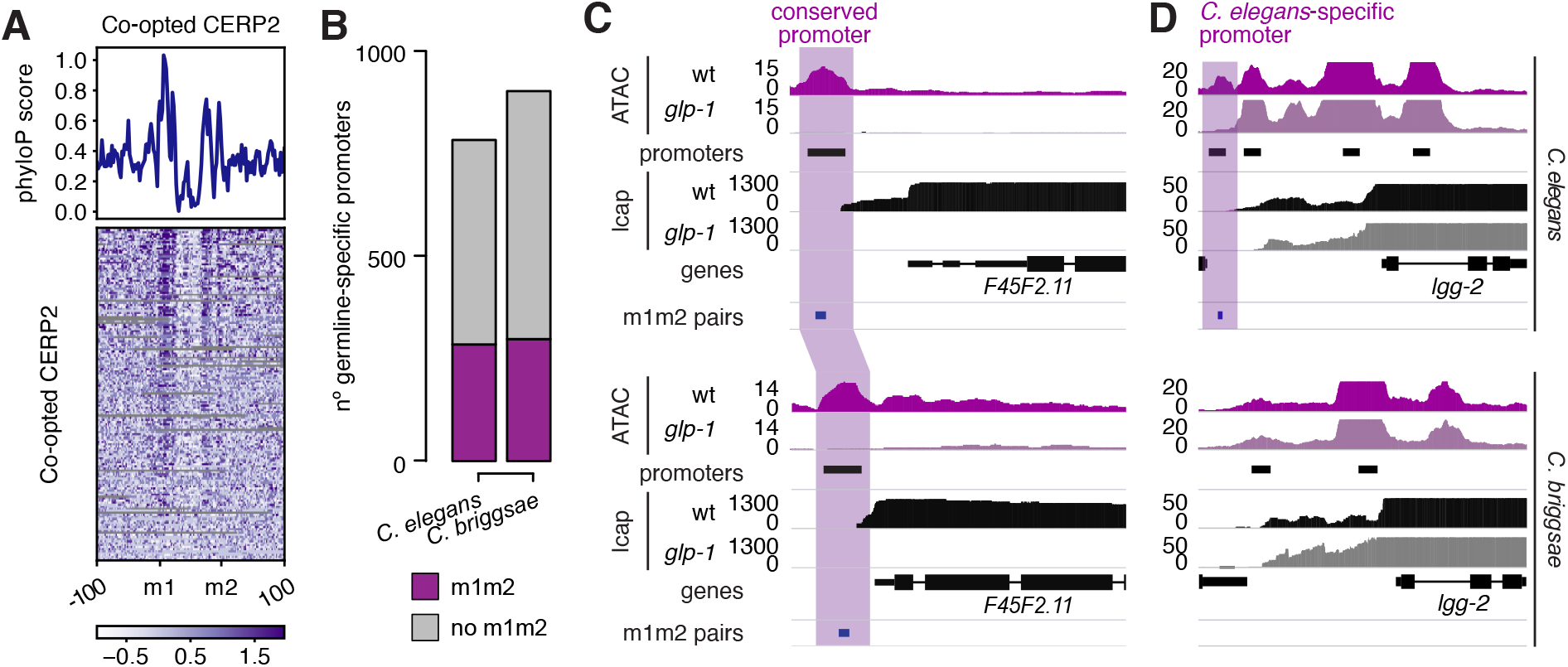
Evolutionary conservation and turnover of co-opted MITEs. (**A**) phyloP score profile (top) and heatmap (bottom) measured at germline-specific promoters associated to a divergent_m1m2 pair in *C. elegans*. Elements not aligned to other species were removed from the heatmap. (**B**) Number of germline-specific promoters annotated in *C. elegans* and *C. briggsae*. (**C,D**) Examples of orthologs with a germline-specific CERP2-derived promoter in *C. elegans* conserved in *C. briggsae* (C) or *C. elegans*-specific (D).

We observed that *C. briggsae* germline-specific promoters, like those in *C. elegans*, are enriched for m1m2 pairs (Fig 4B, Fig S4B). To evaluate the evolutionary conservation of the CERP2 co-opted promoters, we identified 1:1 orthologs associated with a co-opted promoter in *C. elegans* (n=327) or in *C. briggsae* (n=322; Table S1). We found that 53% of the orthologs in each species were regulated by an evolutionary conserved co-opted promoter, and a further 22-27% had some evidence of conservation, indicated either by a promoter with only m1 or m2, or by an m1m2 pair not annotated as a promoter (see Methods). Thus, 53 - 80% of CERP2 co-option events are conserved in *C. elegans* and *C. briggsae* (Fig 4C, S4C and Table 1). The remaining 20-25% of the ortholog pairs had a co-opted promoter in only one species (Fig 4D, S4C and Table 1). This considerable evolutionary turnover could be explained either by the species-specific co-option of new or ancestral MITEs, or by the degeneration of ancestral m1m2 sequences.

Our work provides functional evidence of a large-scale concerted co-option of transposable elements as tissue-specific regulatory sequences. By uncovering hundreds of co-opted promoters preserved by selection for millions of years, we demonstrate that TEs can have a profound impact on the host regulatory landscape. Our identification of this co-option was possible because the promoters still share significant sequence similarity to the MITE elements, whereas the origin of more degenerate or shorter regulatory sequences would be more difficult to trace. Our discovery of widespread co-option of TE sequences as promoters in *Caenorhabditis* supports the possibility that a significant fraction of regulatory sequences in all organisms may originate from transposable elements.

## Materials and Methods

### Strains and growth conditions

*C. elegans* strains were cultured using standard methods (24). A complete list of strains is presented in Table S1.

### Generation of a *C. briggsae glp-1*(*ts*) allele

CRISPR-Cas9 genome editing was used to generate the *C. briggsae glp-1*(*we58*) strain. Injections were performed using gRNA-Cas9 ribonucleoprotein (RNP) complexes preassembled in vitro with in-house made Cas9 protein (25, 26). tracrRNA and crRNAs were purchased from Integrated DNA Technologies and repair templates were Ultramer oligonucleotides from IDT. crRNAs were designed using the online CRISPOR tool (27). We engineered two different mutations in the *C. briggsae glp-1* gene, R955C (GCA −> TCT) and G1036E (GGA −> GAA), to attempt to mimic the *C. elegans* temperature-sensitive e2141 and q231 alleles, respectively. Single mutants did not display germline defects, but each produced some dead eggs at 27°C. We thus generated a double mutant, *glp-1*(*we58*) (carrying both R955C and G1036E). Double mutants were maintained at 16°C and failed to develop a germline when grown from starved L1s at 27°C.

### Generation of *C. briggsae* ATAC-seq and nuclear RNA-seq data

Wild-type AH16 *C. briggsae* or *glp-1*(*we58*) mutants were grown in liquid culture from the starved L1 to the young adult stage using standard S-basal medium with HB101 bacteria (wt at 20C, glp-1 at 27C), frozen in liquid nitrogen, and stored at −80C until use. Nuclei were isolated and ATAC-seq and nuclear RNA-seq libraries generated from wild-type and *glp-1(we58) C. briggsae* young adults as in (14). ATAC-seq and RNA-seq libraries were generated using one million nuclei and for two biological replicates for each *C. briggsae* strain.

### Processing of sequencing data

ChIP-seq data generated in this study, ATAC-seq data from isolated L1 PGCs (GEO accession: GSE100651)(28), and ATAC-seq data from adult germlines (GEO accession: GSE141213)(14) were preprocessed using trim galore (available at https://github.com/FelixKrueger/TrimGalore, version 0.6.4) and mapped using bwa mem (29)(version 0.7.17). Read depth-normalised coverage tracks from mapq10 reads were generated using MACS2 (30)(version 2.1.2; for ATAC-seq data processing, we used the following parameters: --nomodel --extsize 150 --shift −75), converted to bigWig, and replicate pairs were used as input to identify peaks with the yapc software (https://github.com/jurgjn/yapc)(31), with --smoothing-window-width set to 100. Peaks passing an IDR cutoff of 0.00001 (for ChIP-seq) or 0.001 (for ATAC-seq) were used in this study. RNA-seq data were aligned on the genome using STAR (32)(version 2.7.5a) to generate coverage tracks. Gene expression was estimated using kallisto (33)(version 0.46.2).

### Genome annotation

Genome, gene and protein annotations were downloaded from the repositories listed in Table S1. For each protein coding gene, we extracted the genomic and protein sequences of its longest transcript. Repeats from Dfam (34)(release 3.1) were annotated in the *C. elegans* genome using the dfamscan.pl script available on the Dfam website. Repeat coordinates are available in Table S1.

### Identification of germline-specific accessible sites in *C. elegans* and *C. briggsae*

Accessible sites in *C. elegans* and *C. briggsae* were identified using ATAC-seq data generated from wt and *glp-1* mutant strains. Single-end ATAC-seq reads were mapped on the respective genome assembly (WS275 for both *C. elegans* and *C. briggsae*) using bwa-backtrack (35), keeping only reads mapped with high-quality (MAPQ > 10) on fully assembled chromosomes. Coverage-normalised tracks generated using MACS2 were used as input for yapc to identify open chromatin regions. To annotate germline-specific accessible sites in each species, we compared ATAC-seq signals in wt and *glp-1* data using DiffBind (36)(version 2.10.0). We defined sites as germline specific when the *glp-1* vs wt LFC < −2 and the adjusted p-value < 0.01.

### Annotation of germline-specific promoters in *C. elegans* and *C. briggsae*

Germline-specific accessible sites were annotated as promoters in the *C. elegans* or the *C. briggsae* genomes, using a slightly modified version of the annotation pipeline from (31), based on patterns of nuclear RNA-seq data, which identifies regions of transcription elongation, and thus marks the outron regions before trans-splicing. In this work, mapped RNA-seq reads from both replicates of each strain were randomly and evenly distributed in two pseudoreplicates to compensate for lower sequencing depth. Accessible sites were annotated as promoters when a) chromatin-associated RNA-seq signal connected the site to an annotated first exon, allowing gaps in RNA-seq signal of up to 200bp; and b) where a significantly higher RNA-seq signal was present in the regions +75bp to +350bp from the midpoint of an open chromatin region (relative to the downstream gene) compared to the −75 to −350bp sequence.

### Motif enrichment and motif pair annotation

We used the MEME suite (37)(version 5.0.5) to identify motifs enriched in germline-specific promoters in *C. elegans* or *C. briggsae* (enrichment compared to non-GL-specific promoters, MEME-ChIP parameters used: -meme-nmotifs 6 -meme-minw 5 -meme-maxw 20).

Enriched motifs were mapped on the genomes of different species with FIMO (P < 0.0005). We annotated all occurrences of m1 and m2 motifs separated by 10-30bp - a range including the most frequently observed m1-m2 spacings - as m1m2 motif pairs, and distinguished them based on the relative motif orientation into 4 arrangements: convergent_m1m2, divergent_m1m2, tandem_m1m2 and tandem_m2m1 (Table S1).

### Assessment of CERP2 and CELE2 derived promoter activity

Transgenes containing the annotated CERP2-associated promoter of *C16A11.4* or the CELE2-associated promoter of *fat-1* upstream of *his-58::gfp::tbb-2* 3’UTR were generated using mosSCI (38). Wild-type and mutant versions in which motifs were scrambled were generated. Promoter sequences used are given in Table S1.

Synthesised promoter sequences were ordered as plasmids containing att sites for Gateway cloning from GenScript, and reporter transgenes constructed using three-site Gateway cloning (Invitrogen), using vector pCFJ150, which targets Mos site Mos1(ttTi5605) on chromosome II (38), the promoter to be tested in site one, his-58 in site two (plasmid pJA357), and gfp-tbb-2 3’UTR in site three (pJA256) (39). GFP signal was assessed using a Zeiss Axioplan microscope equipped with wide-field fluorescence microscopy. At least 20 individuals were scored per strain.

### *T05F1.2* promoter mutation

We used CRISPR-Cas9 to scramble the m1 and m2 sequences in the endogenous CERP2-associated promoter of *T05F1.2. T05F1.2* expression in the wild-type and mutant strain (*we59*) was quantified by qPCR using two different sets of primers and compared to *cdc-42* expression. Primer sequences used are available in Table S1.

### Association of co-opted promoters with TF binding sites

ChIP-seq data from 283 *C. elegans* TFs were downloaded as aggregated peaks from the modERN website (https://epic.gs.washington.edu/modERN/)(15), and from these we extracted only data from 73 factors which were generated from young adult animals. We further included data from a single HIM-17 ChIP-seq replicate (3916_SDQ0801_HIM17_FEM2_AD_r1) available in modENCODE (40) but not included in modERN. The HIM-17 ChIP-seq reads were mapped on the ce11 genome using bwa-mem, and peaks were called using mapq10 reads with MACS2. For each factor we compared the ratio of peaks overlapping germline-specific co-opted and non-co-opted promoters.

### HIM-17 ChIP-seq

HIM-17 ChIP-seq libraries were prepared from two biological replicates following the protocol described in (41). Heatmaps of HIM-17 ChIP-seq profiles, and its association to CERP2 or CELE2 repeats, m1m2 pairs, and regulatory elements were generated using the computeMatrix and plotHeatmap functions from the deepTools2 suite (42)(version 3.4.3).

### Testing requirement for m1m2 motifs in HIM-17 chromatin association

To test if HIM-17 requires motifs m1 or m2 for chromatin association at a co-opted promoter, three variants of the transgene driven by the CERP2-derived *C16A11.4* promoter were generated using MosSCI: scrambled m1, scrambled m2, or scrambled m1 and m2. ChIP-qPCR was performed for HIM-17, testing enrichment for the transgene promoter, for the co-opted *ztf-15* promoter as a positive control, and for two negative control loci showing no ChIP-seq enrichment for either factor. Experiments were done on three technical replicates from two biological replicates.

### *him-17* gene expression analysis

For each of two replicates, approximately 100 wild-type and *him-17*(*me24*) (m+z-) young adults grown at 20°C from the starved L1 stage. *him-17*(*me24*) were derived from *him-17*(*me24*)/*tmC12* [*tmIs1194*] mothers. Total RNA was extracted using Trizol. poly A was isolated using the NEBNext Poly(A) mRNA Isolation kit and libraries were prepared using the NEBNext Ultra Directional RNA Library Prep Kit (E7760S).

DESeq2 (43)(version 1.22.1) was used to identify significantly upregulated (LFC > 0, p.adj < 0.001) or downregulated (LFC < 0, p.adj < 0.001) genes in him-17 mutants compared to wild-type. GO enrichment analysis on differentially expressed genes was performed with clusterProfile (44). Direct targets were defined as differentially expressed genes that have a HIM-17 ChIP-seq peak on their promoter.

### Annotation of CERP2 and CELE2 in different species

We extracted sequences from all CERP2 and CELE2 elements in *C. elegans* to refine HMM models of these repeats using the HMMER3 suite (http://hmmer.org/). Fasta sequences of all repeats from each family were aligned against the CERP2 or CELE2 Dfam HMM using hmmalign (with parameter --trim). The resulting alignment was used to define new HMMs using hmmbuild. The HMMs were then used to annotate CERP2 and CELE2 repeats in nematodes with chromosome-level genome annotations (Table S1) using nhmmer and requiring a minimal E-value of 0.001.

### HIM-17 evolution and structure

HIM-17 orthologs were identified using BLASTP (E-value < 0.00001) on the protein annotation from a number of nematode species. To test the *him-17* sequence for positive selection, HIM-17 orthologs were aligned using MAFFT (45) with the L-INS-i method, then the output alignment was used to guide a codon-based alignment using PAL2NAL (46). The resulting alignment was used to test for positive selection acting on the common *Caenorhabditis* branch using the branch-site test (47) implemented in codeml from the PAML package (48). THAP domains in HIM-17 orthologs were annotated with hmmsearch using the THAP profile HMMs from the Pfam database (49).

### Analysis of co-opted promoters conservation

Sequence conservation of m1m2 pairs located in CERP2-derived germline-specific promoters was assessed using phyloP scores from 26 nematodes (phyloP26way from the ce11 release) available from the UCSC genome browser.

To evaluate the conservation of individual CERP2 promoters in *C. elegans* and *C. briggsae*, we extracted all 1-to-1 orthologs (obtained from Wormbase) regulated by a co-opted promoter in at least one species, i.e. associated to at least one promoter containing an m1m2 pair in divergent orientation and spaced by 12 to 16 bp (CERP2-like arrangement). Co-opted promoters were defined as conserved when both orthologs were associated with a co-opted promoter. We considered co-opted promoters as potentially conserved when the ortholog in the other species was either a) associated with a promoter containing at least m1 or m2; b) when an m1m2 pair was located in the putative promoter region (−1000bp/+200bp) of the orthologs’ TSS but was not in an annotated promoter. When none of the criteria were met, we defined the co-opted promoter as species-specific.

## Data availability

ATAC-seq, ChIP-seq and RNA-seq data generated during this study are available at NCBI Gene Expression Omnibus (GEO) under accession code GSE192540. The code used for the analysis is available on GitHub (https://github.com/fcarelli/glre).

## Acknowledgments

The authors thank current and former members of the Ahringer lab for helpful discussions and Jürgen Jänes and Andrea Frapporti for experimental assistance.

## Author contributions

Conceptualization: FNC, JA; Methodology: FNC, JA; Investigation: FNC, CC, YD, AA; Visualization: FNC, JA; Funding acquisition: JA; Project administration: JA; Supervision: JA; Writing – original draft: FNC, JA; Writing – review & editing: FNC, JA, AD.

---

**Fig. S1.**
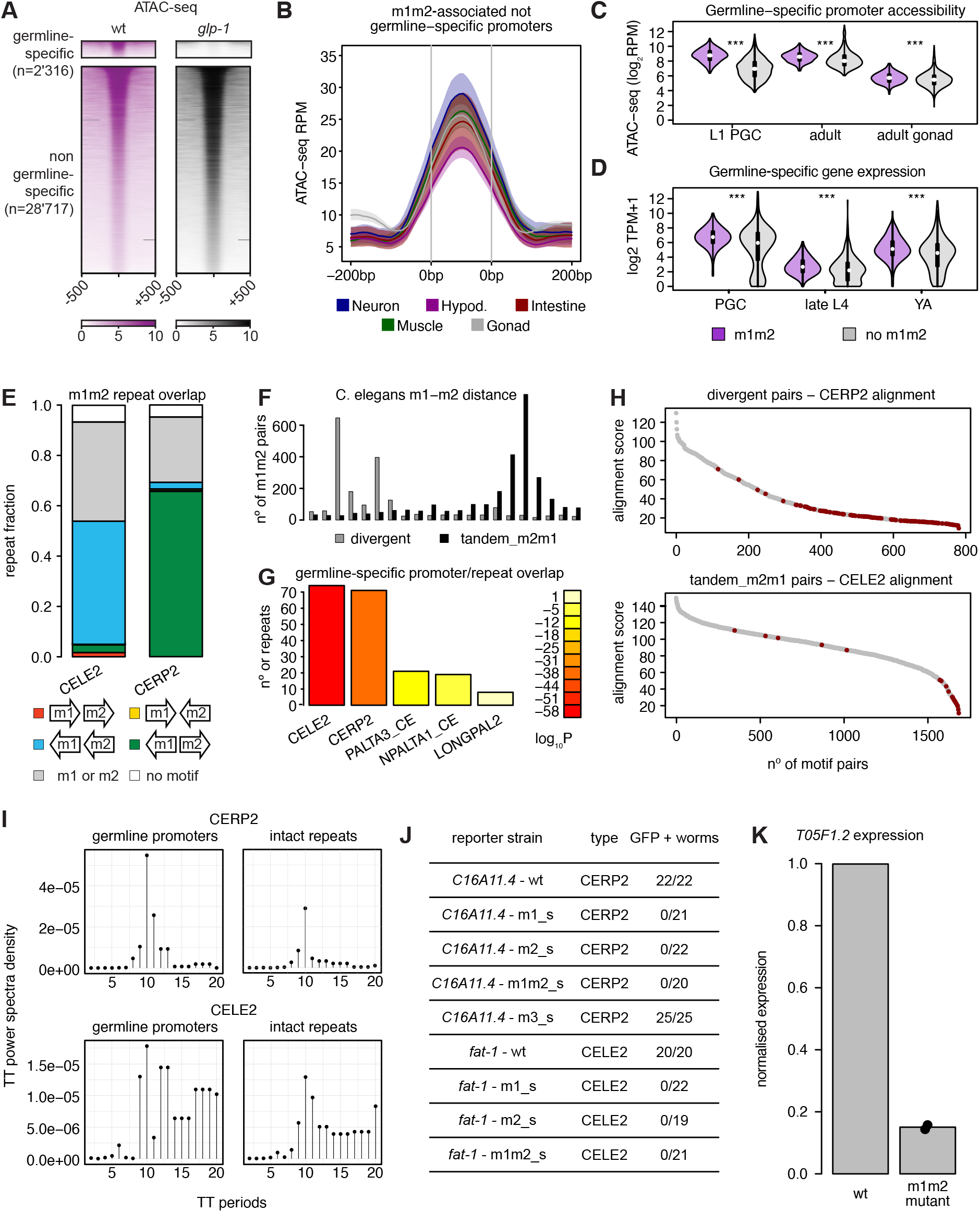
TE enrichment at germline-specific elements in *C. elegans*. (**A**) ATAC seq signal at *C. elegans* open chromatin regions. (**B**) ATAC-seq coverage from individual tissues (from Serizay et al.) over non-germline-specific promoters associated with an m1m2 pair. (**C**) ATAC-seq coverage at germline-specific promoters. (**D**) Expression levels of genes regulated by a unique germline-specific promoter. (**E**) Fraction of CERP2 and CELE2 elements overlapping m1m2 pairs in any arrangement. (**F**) Spacing between m1 and m2 motifs in divergent and tandem_m2m1 pairs. (**G**) Enrichment of repeats families in germline-specific vs non germline-specific promoters. Only families with a significant enrichment above 1 were depicted. (**H**) Alignment score of m1m2 pairs in divergent or tandem_m2m1 arrangement to the CERP2 or CELE2 repeat model, respectively. Motifs in promoters are depicted in red. (**I**) Power Spectral Densities of TT dinucleotides measured downstream of intact and germline-specific promoter-associated divergent_m1m2 and tandem_m2m1 pairs. (**J**) Fraction of young adult hermaphrodites carrying indicated transgenes with GFP expression in the germ line. (**K**) Fold-change (measured by qPCR) in expression of the *T05F1.2* gene after mutating its promoter compared to wt.

**Fig. S2.**
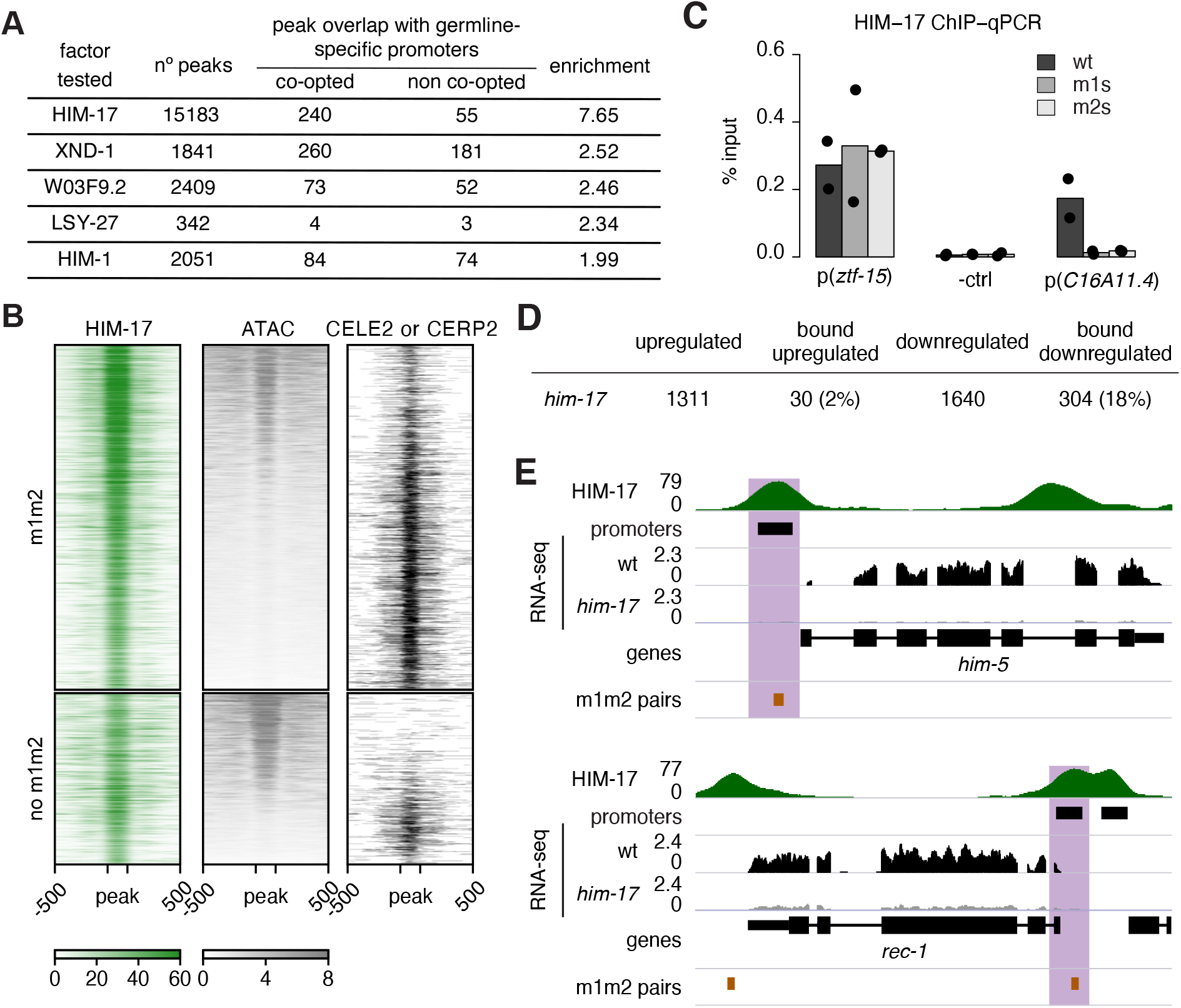
HIM-17 and XND-1 bind co-opted and inactive MITEs. (**A**) Summary statistics of top overlaps between co-opted and non-co-opted germline-specific promoters and the modERN/modENCODE peaks set. (**B**) HIM-17, ATAC-seq and CERP2 or CELE2 enrichment over HIM-17 peaks. Top, peaks overlapping an annotated m1m2 pair (n=2364); bottom, peaks without an annotated m1m2 pair (n=1175). (**C**) ChIP-qPCR enrichment of HIM-17 as % of input in strains containing the wt, m1 and m2 scrambled versions of the *C16A11.4* promoter integrated in the chrII MosSCI site (see Methods). Signal at endogenous co-opted promoter *p*(*ztf-15*) is shown as a positive control. (**D**) number of upregulated (up) and downregulated genes (down) in *him-17* mutants. When the factor (in wt) overlapped any of the genes differentially expressed in the corresponding mutant, the gene was considered a direct target. Percentages: fraction of direct targets over all DE genes. (**E**) HIM-17 binding profile and RNA-seq profiles in wt and *him-17* mutants at the *him-5* locus.

**Fig. S3.**
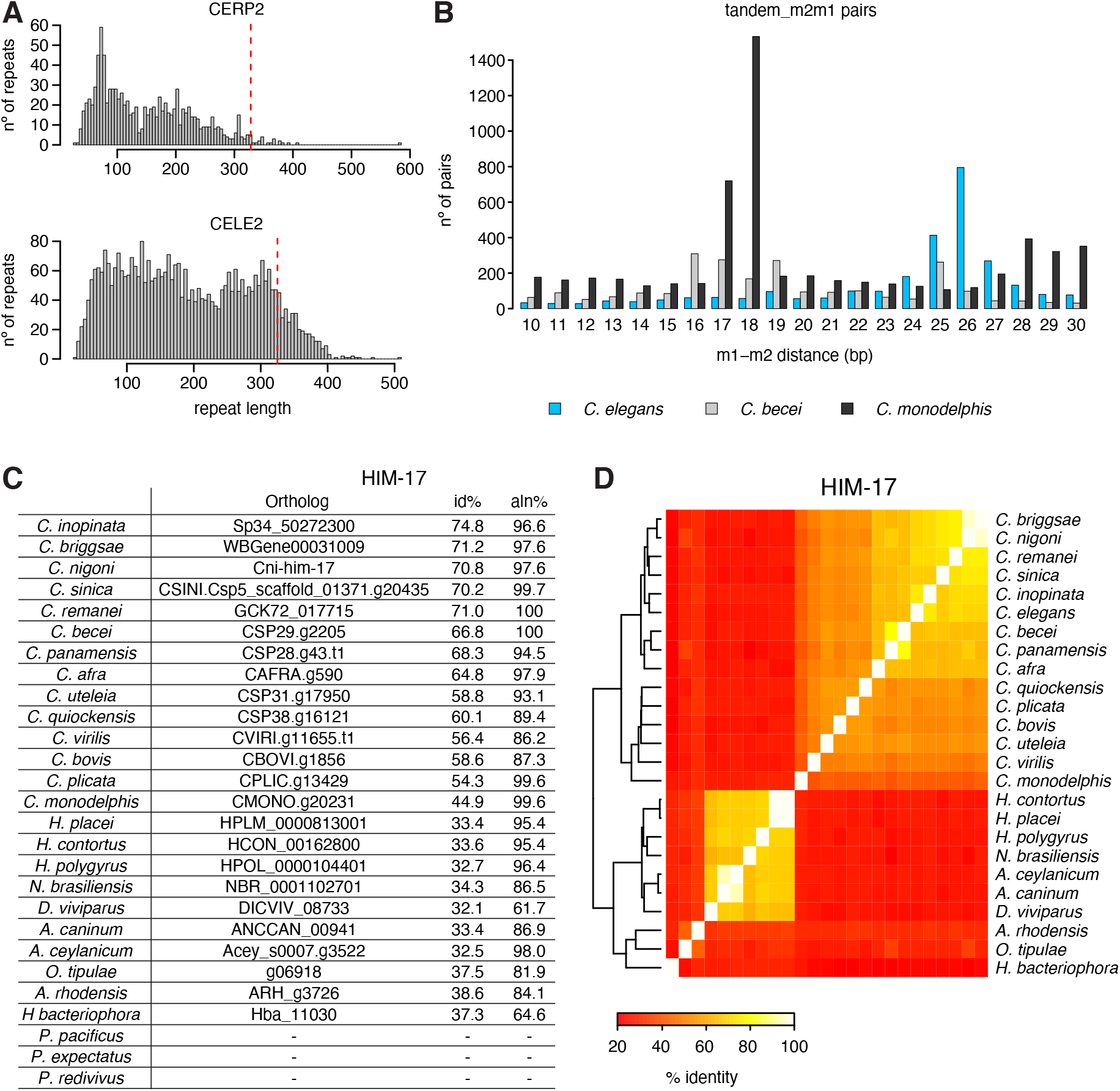
Evolution of m1m2 pairs and their binding factors in nematodes. (**A**) Length of annotated CERP2 and CELE2 elements in *C. elegans*; red dashed line indicates length of consensus repeat sequence. (**B**) Spacing between m1 and m2 motifs in tandem_m2m1 in a subset of species. (**C**) HIM-17 orthologs in different nematodes, with % identity and % of alignment length with the corresponding *C. elegans* protein. (**D**) pairwise % identity across HIM-17 orthologs.

**Fig. S4.**
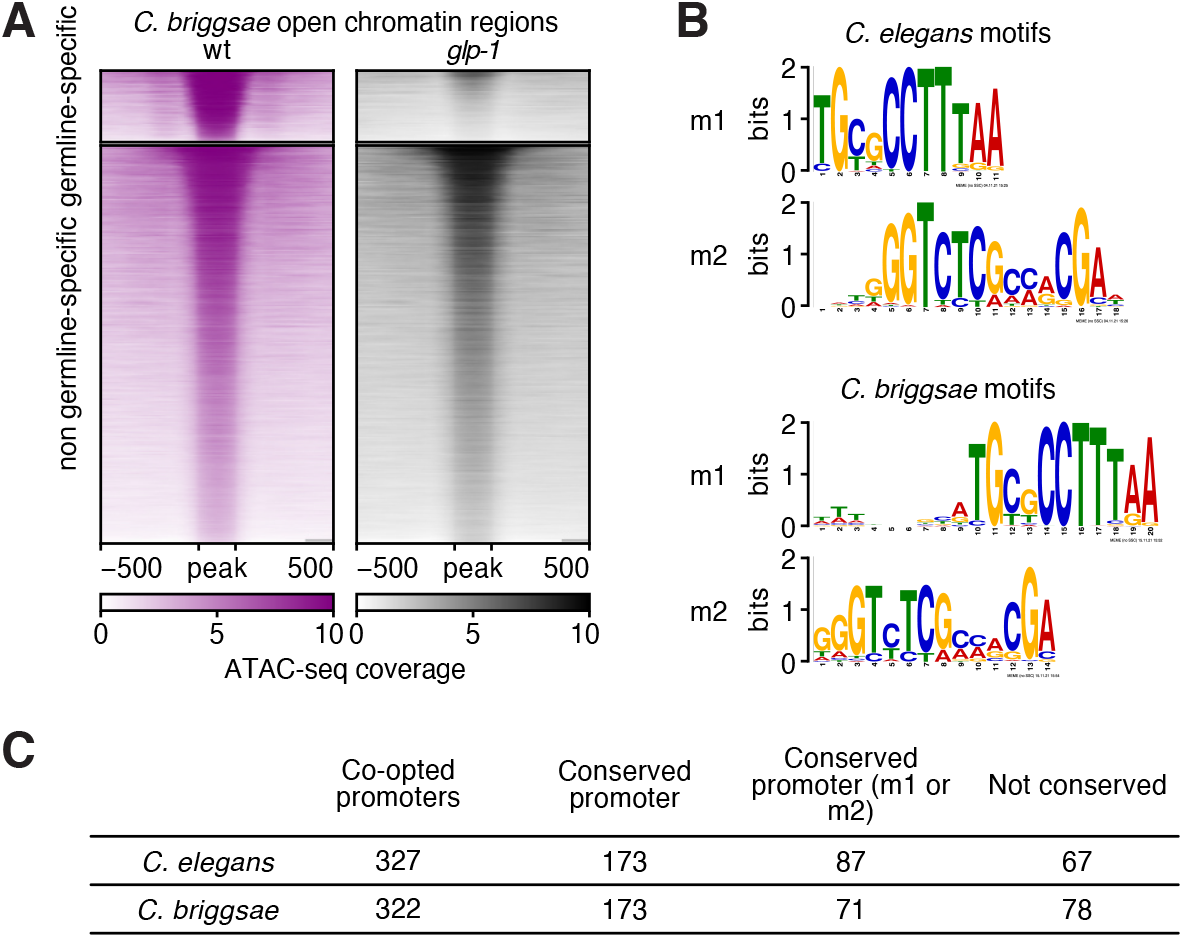
Evolutionary conservation and turnover of co-opted MITEs. (**A**) ATAC-seq signal (in RPM) from wild-type and *Cbr-glp-1* mutant over *C. briggsae* open chromatin regions. (**B**) comparison of m1 and m2 motif logos in *C. elegans* and *C. briggsae*. (**C**) summary of CERP2 promoter conservation between *C. elegans* and *C. briggsae* orthologs.

## References

1. R. K. Bradley, X.-Y. Li, C. Trapnell, S. Davidson, L. Pachter, H. C. Chu, L. A. Tonkin, M. D. Biggin, M. B. Eisen, Binding site turnover produces pervasive quantitative changes in transcription factor binding between closely related Drosophila species. PLoS Biol. 8, e1000343 (2010).

2. D. Villar, C. Berthelot, S. Aldridge, T. F. Rayner, M. Lukk, M. Pignatelli, T. J. Park, R. Deaville, J. T. Erichsen, A. J. Jasinska, J. M. A. Turner, M. F. Bertelsen, E. P. Murchison, P. Flicek, D. T. Odom, Enhancer evolution across 20 mammalian species. Cell. 160, 554–566 (2015).

3. R. S. Young, Y. Hayashizaki, R. Andersson, A. Sandelin, H. Kawaji, M. Itoh, T. Lassmann, P. Carninci, FANTOM Consortium, W. A. Bickmore, A. R. Forrest, M. S. Taylor, The frequent evolutionary birth and death of functional promoters in mouse and human. Genome Res. 25, 1546–1557 (2015).

4. E. B. Chuong, N. C. Elde, C. Feschotte, Regulatory activities of transposable elements: from conflicts to benefits. Nat. Rev. Genet. 18, 71–86 (2017).

5. R. L. Cosby, N.-C. Chang, C. Feschotte, Host-transposon interactions: conflict, cooperation, and cooption. Genes Dev. 33, 1098–1116 (2019).

6. E. B. Chuong, M. A. K. Rumi, M. J. Soares, J. C. Baker, Endogenous retroviruses function as species-specific enhancer elements in the placenta. Nat. Genet. 45, 325–329 (2013).

7. V. J. Lynch, M. C. Nnamani, A. Kapusta, K. Brayer, S. L. Plaza, E. C. Mazur, D. Emera, S. Z. Sheikh, F. Grützner, S. Bauersachs, A. Graf, S. L. Young, J. D. Lieb, F. J. DeMayo, C. Feschotte, G. P. Wagner, Ancient transposable elements transformed the uterine regulatory landscape and transcriptome during the evolution of mammalian pregnancy. Cell Rep. 10, 551–561 (2015).

8. E. B. Chuong, N. C. Elde, C. Feschotte, Regulatory evolution of innate immunity through co-option of endogenous retroviruses. Science. 351, 1083–1087 (2016).

9. A. Sakashita, S. Maezawa, K. Takahashi, K. G. Alavattam, M. Yukawa, Y.-C. Hu, S. Kojima, N. F. Parrish, A. Barski, M. Pavlicev, S. H. Namekawa, Endogenous retroviruses drive species-specific germline transcriptomes in mammals. Nat. Struct. Mol. Biol. 27, 967–977 (2020).

10. J. Austin, J. Kimble, glp-1 is required in the germ line for regulation of the decision between mitosis and meiosis in C. elegans. Cell. 51, 589–599 (1987).

11. C. Linhart, Y. Halperin, A. Darom, S. Kidron, L. Broday, R. Shamir, A novel candidate cis-regulatory motif pair in the promoters of germline and oogenesis genes in C. elegans. Genome Res. 22, 76–83 (2012).

12. C. Feschotte, C. Mouchès, Evidence that a family of miniature inverted-repeat transposable elements (MITEs) from the Arabidopsis thaliana genome has arisen from a pogo-like DNA transposon. Mol. Biol. Evol. 17, 730–737 (2000).

13. C. Feschotte, E. J. Pritham, DNA transposons and the evolution of eukaryotic genomes. Annu. Rev. Genet. 41, 331–368 (2007).

14. J. Serizay, Y. Dong, J. Jänes, M. Chesney, C. Cerrato, J. Ahringer, Distinctive regulatory architectures of germline-active and somatic genes in C. elegans. Genome Res. 30, 1752–1765 (2020).

15. M. M. Kudron, A. Victorsen, L. Gevirtzman, L. W. Hillier, W. W. Fisher, D. Vafeados, M. Kirkey, A. S. Hammonds, J. Gersch, H. Ammouri, M. L. Wall, J. Moran, D. Steffen, M. Szynkarek, S. Seabrook-Sturgis, N. Jameel, M. Kadaba, J. Patton, R. Terrell, M. Corson, T. J. Durham, S. Park, S. Samanta, M. Han, J. Xu, K.-K. Yan, S. E. Celniker, K. P. White, L. Ma, M. Gerstein, V. Reinke, R. H. Waterston, The ModERN Resource: Genome-Wide Binding Profiles for Hundreds of Drosophila and Caenorhabditis elegans Transcription Factors. Genetics. 208, 937–949 (2018).

16. K. C. Reddy, A. M. Villeneuve, C. elegans HIM-17 links chromatin modification and competence for initiation of meiotic recombination. Cell. 118, 439–452 (2004).

17. P. M. Meneely, O. L. McGovern, F. I. Heinis, J. L. Yanowitz, Crossover distribution and frequency are regulated by him-5 in Caenorhabditis elegans. Genetics. 190, 1251–1266 (2012).

18. M. Roussigne, S. Kossida, A.-C. Lavigne, T. Clouaire, V. Ecochard, A. Glories, F. Amalric, J.-P. Girard, The THAP domain: a novel protein motif with similarity to the DNA-binding domain of P element transposase. Trends Biochem. Sci. 28, 66–69 (2003).

19. G. Chung, A. M. Rose, M. I. R. Petalcorin, J. S. Martin, Z. Kessler, L. Sanchez-Pulido, C. P. Ponting, J. L. Yanowitz, S. J. Boulton, REC-1 and HIM-5 distribute meiotic crossovers and function redundantly in meiotic double-strand break formation in Caenorhabditis elegans. Genes Dev. 29, 1969–1979 (2015).

20. S. Nadarajan, E. Altendorfer, T. T. Saito, M. Martinez-Garcia, M. P. Colaiácovo, HIM-17 regulates the position of recombination events and GSP-1/2 localization to establish short arm identity on bivalents in meiosis. Proc. Natl. Acad. Sci. U. S. A. 118 (2021), doi:10.1073/pnas.2016363118.

21. J. B. Bessler, K. C. Reddy, M. Hayashi, J. Hodgkin, A. M. Villeneuve, A role for Caenorhabditis elegans chromatin-associated protein HIM-17 in the proliferation vs. meiotic entry decision. Genetics. 175, 2029–2037 (2007).

22. X. She, X. Xu, A. Fedotov, W. G. Kelly, E. M. Maine, Regulation of heterochromatin assembly on unpaired chromosomes during Caenorhabditis elegans meiosis by components of a small RNA-mediated pathway. PLoS Genet. 5, e1000624 (2009).

23. A. D. Cutter, Divergence times in Caenorhabditis and Drosophila inferred from direct estimates of the neutral mutation rate. Mol. Biol. Evol. 25, 778–786 (2008).

24. S. Brenner, The genetics of Caenorhabditis elegans. Genetics. 77, 71–94 (1974).

25. A. Paix, A. Folkmann, D. Rasoloson, G. Seydoux, High Efficiency, Homology-Directed Genome Editing in Caenorhabditis elegans Using CRISPR-Cas9 Ribonucleoprotein Complexes. Genetics. 201, 47–54 (2015).

26. A. Paix, A. Folkmann, D. H. Goldman, H. Kulaga, M. J. Grzelak, D. Rasoloson, S. Paidemarry, R. Green, R. R. Reed, G. Seydoux, Precision genome editing using synthesis-dependent repair of Cas9-induced DNA breaks. Proc. Natl. Acad. Sci. U. S. A. 114, E10745–E10754 (2017).

27. M. Haeussler, K. Schönig, H. Eckert, A. Eschstruth, J. Mianné, J.-B. Renaud, S. Schneider-Maunoury, A. Shkumatava, L. Teboul, J. Kent, J.-S. Joly, J.-P. Concordet, Evaluation of off-target and on-target scoring algorithms and integration into the guide RNA selection tool CRISPOR. Genome Biol. 17, 148 (2016).

28. C.-Y. S. Lee, T. Lu, G. Seydoux, Nanos promotes epigenetic reprograming of the germline by down-regulation of the THAP transcription factor LIN-15B. Elife. 6 (2017), doi:10.7554/eLife.30201.

29. H. Li, Aligning sequence reads, clone sequences and assembly contigs with BWA-MEM. arXiv [q-bio.GN] (2013), (available at http://arxiv.org/abs/1303.3997).

30. Y. Zhang, T. Liu, C. A. Meyer, J. Eeckhoute, D. S. Johnson, B. E. Bernstein, C. Nusbaum, R. M. Myers, M. Brown, W. Li, X. S. Liu, Model-based analysis of ChIP-Seq (MACS). Genome Biol. 9, R137 (2008).

31. J. Jänes, Y. Dong, M. Schoof, J. Serizay, A. Appert, C. Cerrato, C. Woodbury, R. Chen, C. Gemma, N. Huang, D. Kissiov, P. Stempor, A. Steward, E. Zeiser, S. Sauer, J. Ahringer, Chromatin accessibility dynamics across C. elegans development and ageing. Elife. 7 (2018), doi:10.7554/eLife.37344.

32. A. Dobin, C. A. Davis, F. Schlesinger, J. Drenkow, C. Zaleski, S. Jha, P. Batut, M. Chaisson, T. R. Gingeras, STAR: ultrafast universal RNA-seq aligner. Bioinformatics. 29, 15–21 (2013).

33. N. L. Bray, H. Pimentel, P. Melsted, L. Pachter, Near-optimal probabilistic RNA-seq quantification. Nat. Biotechnol. 34, 525–527 (2016).

34. R. Hubley, R. D. Finn, J. Clements, S. R. Eddy, T. A. Jones, W. Bao, A. F. A. Smit, T. J. Wheeler, The Dfam database of repetitive DNA families. Nucleic Acids Res. 44, D81–9 (2016).

35. H. Li, R. Durbin, Fast and accurate long-read alignment with Burrows-Wheeler transform. Bioinformatics. 26, 589–595 (2010).

36. C. S. Ross-Innes, R. Stark, A. E. Teschendorff, K. A. Holmes, H. R. Ali, M. J. Dunning, G. D. Brown, O. Gojis, I. O. Ellis, A. R. Green, S. Ali, S.-F. Chin, C. Palmieri, C. Caldas, J. S. Carroll, Differential oestrogen receptor binding is associated with clinical outcome in breast cancer. Nature. 481, 389–393 (2012).

37. T. L. Bailey, M. Boden, F. A. Buske, M. Frith, C. E. Grant, L. Clementi, J. Ren, W. W. Li, W. S. Noble, MEME SUITE: tools for motif discovery and searching. Nucleic Acids Res. 37, W202–8 (2009).

38. C. Frøkjaer-Jensen, M. W. Davis, C. E. Hopkins, B. J. Newman, J. M. Thummel, S.-P. Olesen, M. Grunnet, E. M. Jorgensen, Single-copy insertion of transgenes in Caenorhabditis elegans. Nat. Genet. 40, 1375–1383 (2008).

39. E. Zeiser, C. Frøkjær-Jensen, E. Jorgensen, J. Ahringer, MosSCI and gateway compatible plasmid toolkit for constitutive and inducible expression of transgenes in the C. elegans germline. PLoS One. 6, e20082 (2011).

40. M. B. Gerstein, Z. J. Lu, E. L. Van Nostrand, C. Cheng, B. I. Arshinoff, T. Liu, K. Y. Yip, R. Robilotto, A. Rechtsteiner, K. Ikegami, P. Alves, A. Chateigner, M. Perry, M. Morris, R. K. Auerbach, X. Feng, J. Leng, A. Vielle, W. Niu, K. Rhrissorrakrai, A. Agarwal, R. P. Alexander, G. Barber, C. M. Brdlik, J. Brennan, J. J. Brouillet, A. Carr, M.-S. Cheung, H. Clawson, S. Contrino, L. O. Dannenberg, A. F. Dernburg, A. Desai, L. Dick, A. C. Dosé, J. Du, T. Egelhofer, S. Ercan, G. Euskirchen, B. Ewing, E. A. Feingold, R. Gassmann, P. J. Good, P. Green, F. Gullier, M. Gutwein, M. S. Guyer, L. Habegger, T. Han, J. G. Henikoff, S. R. Henz, A. Hinrichs, H. Holster, T. Hyman, A. L. Iniguez, J. Janette, M. Jensen, M. Kato, W. J. Kent, E. Kephart, V. Khivansara, E. Khurana, J. K. Kim, P. Kolasinska-Zwierz, E. C. Lai, I. Latorre, A. Leahey, S. Lewis, P. Lloyd, L. Lochovsky, R. F. Lowdon, Y. Lubling, R. Lyne, M. MacCoss, S. D. Mackowiak, M. Mangone, S. McKay, D. Mecenas, G. Merrihew, D. M. Miller 3rd, A. Muroyama, J. I. Murray, S.-L. Ooi, H. Pham, T. Phippen, E. A. Preston, N. Rajewsky, G. Rätsch, H. Rosenbaum, J. Rozowsky, K. Rutherford, P. Ruzanov, M. Sarov, R. Sasidharan, A. Sboner, P. Scheid, E. Segal, H. Shin, C. Shou, F. J. Slack, C. Slightam, R. Smith, W. C. Spencer, E. O. Stinson, S. Taing, T. Takasaki, D. Vafeados, K. Voronina, G. Wang, N. L. Washington, C. M. Whittle, B. Wu, K.-K. Yan, G. Zeller, Z. Zha, M. Zhong, X. Zhou, modENCODE Consortium, J. Ahringer, S. Strome, K. C. Gunsalus, G. Micklem, X. S. Liu, V. Reinke, S. K. Kim, L. W. Hillier, S. Henikoff, F. Piano, M. Snyder, L. Stein, J. D. Lieb, R. H. Waterston, Integrative analysis of the Caenorhabditis elegans genome by the modENCODE project. Science. 330, 1775–1787 (2010).

41. A. N. McMurchy, P. Stempor, T. Gaarenstroom, B. Wysolmerski, Y. Dong, D. Aussianikava, A. Appert, N. Huang, P. Kolasinska-Zwierz, A. Sapetschnig, E. A. Miska, J. Ahringer, A team of heterochromatin factors collaborates with small RNA pathways to combat repetitive elements and germline stress. Elife. 6 (2017), doi:10.7554/eLife.21666.

42. F. Ramírez, D. P. Ryan, B. Grüning, V. Bhardwaj, F. Kilpert, A. S. Richter, S. Heyne, F. Dündar, T. Manke, deepTools2: a next generation web server for deep-sequencing data analysis. Nucleic Acids Res. 44, W160–5 (2016).

43. M. I. Love, W. Huber, S. Anders, Moderated estimation of fold change and dispersion for RNA-seq data with DESeq2. Genome Biol. 15, 550 (2014).

44. G. Yu, L.-G. Wang, Y. Han, Q.-Y. He, clusterProfiler: an R package for comparing biological themes among gene clusters. OMICS. 16, 284–287 (2012).

45. K. Katoh, K.-I. Kuma, H. Toh, T. Miyata, MAFFT version 5: improvement in accuracy of multiple sequence alignment. Nucleic Acids Res. 33, 511–518 (2005).

46. M. Suyama, D. Torrents, P. Bork, PAL2NAL: robust conversion of protein sequence alignments into the corresponding codon alignments. Nucleic Acids Res. 34, W609–12 (2006).

47. J. Zhang, R. Nielsen, Z. Yang, Evaluation of an improved branch-site likelihood method for detecting positive selection at the molecular level. Mol. Biol. Evol. 22, 2472–2479 (2005).

48. Z. Yang, PAML 4: phylogenetic analysis by maximum likelihood. Mol. Biol. Evol. 24, 1586–1591 (2007).

49. J. Mistry, S. Chuguransky, L. Williams, M. Qureshi, G. A. Salazar, E. L. L. Sonnhammer, S. C. E. Tosatto, L. Paladin, S. Raj, L. J. Richardson, R. D. Finn, A. Bateman, Pfam: The protein families database in 2021. Nucleic Acids Res. 49, D412–D419 (2021).

